# An Engineered Cas-Transposon System for Programmable and Precise DNA Transpositions

**DOI:** 10.1101/654996

**Authors:** Sway P. Chen, Harris H. Wang

## Abstract

Efficient targeted insertion of heterologous DNA into a genome remains a challenge in genome engineering. Recombinases that can introduce kilobase-sized DNA constructs require pre-existing recombination sites to be present in the genome and are difficult to reprogram to other loci. Genome insertion using current CRISPR-Cas methods relies on host DNA repair machinery, which is generally inefficient. Here, we describe a Cas-Transposon (CasTn) system for genomic insertions that uses a transposase fused to a catalytically-dead dCas9 nuclease to mediate programmable, site-specific transposition. CasTn combines the power of the Himar1 transposase, which inserts multi-kb DNA transposons into TA dinucleotides by a cut-and-paste mechanism, and the targeting capability of Cas9, which uses guide-RNAs to bind to specific DNA sequences. Using *in vitro* assays, we demonstrated that Himar-dCas9 proteins increased the frequency of transposon insertions at a single targeted TA dinucleotide by >300-fold compared to an untargeted transposase, and that site-specific transposition is dependent on target choice while robust to log-fold variations in protein and DNA concentrations. We then showed that Himar-dCas9 mediates site-specific transposition into a target plasmid in *E. coli*. This work provides CasTn as a new method for host-independent, programmable, targeted DNA insertions to expand the genomic engineering toolbox.

## INTRODUCTION

Genome engineering relies on molecular tools for targeted and specific modification of a genome to introduce insertions, deletions, and substitutions. While numerous advances have emerged over the last decade to enable programmable editing and deletion of bacterial and eukaryotic genomes^1^, targeted genomic insertion has been a long-outstanding challenge. Integration of desired heterologous DNA into the genome needs to be precise, programmable, and efficient, three key parameters of any genome integration methodology. Currently available genome integration tools are limited by one or more of these factors. Recombinases such as Flp^2^ and Cre^3^ that mediate recombination at defined recognition sequences to integrate heterologous DNA have limited programmability, since their DNA-recognition domains are challenging to reengineer to target other DNA motifs^4, 5^. Site-specific nucleases, such as Cas9^6, 7^, zinc-finger nucleases (ZFNs)^8^, and transcription activator-like effector nucleases (TALENs)^9^, can be programmed to generate double-strand DNA breaks that are then repaired to incorporate a template DNA. However, this process relies on host homology-directed repair machinery, which is variable and often inefficient, especially as the size of the DNA insertion increases^10^.

Transposable elements are widespread natural selfish genetic systems capable of integrating large pieces of DNA into both prokaryotic and eukaryotic genomes. Amongst various transposable elements described^11, 12^, the Himar1 transposon from the horn fly *Haematobia irritans*^13^ has been coopted as a popular tool for insertional mutagenesis. The Himar1 transposon is mobilized by the Himar1 transposase, which like other Tc1/*mariner*-family transposases, functions as a homodimer to bind the transposon DNA at the flanking inverted repeat sequences, excise the transposon, and paste it into a random TA dinucleotide on a target DNA^13-16^. Himar1 transposition requires no host factors for transposition and functions *in vitro*^13^, in bacteria^17^, and in mammalian cells^18^, and is capable of inserting transposons over 7 kb in size^19^. A hyperactive mutant of the transposase, Himar1C9, which contains two amino acid substitutions and increases transposition efficiency by 50-fold^20^, has enabled the generation of genome-wide transposon insertion mutant libraries for genetic screens in diverse microbes^21-23^. However, because Himar1 transposons are inserted randomly into TA dinucleotide sites, their utility in targeted genome insertion applications has been limited thus far.

One approach to potentially increase the specificity of otherwise random transposon insertions is to increase the affinity of the transposase to specific DNA motifs. Indeed, previous studies have described fusing transposases to a DNA-binding protein (DBP) domain to increase targeting of transposon insertions to specific genetic loci. Fusing the Gal4 DNA-binding protein to Mos1 (a Tc1/*mariner* family member) and piggyBac transposases altered the distribution of integration sites to near Gal4 recognition sites in plasmid-based assays in mosquito embryos^24^. Fusion of DNA-binding zinc-finger or transcription activator-like (TAL) effector proteins to piggyBac enabled integration into specified genomic loci in human cells^25-27^. ISY100 transposase (also a Tc1/*mariner* family member) has been fused to a Zif268 Zinc-finger domain to increase specificity of transposon insertions to DNA adjacent to Zif268 binding sites^28^. While these studies demonstrated the feasibility of the transposase-DBP fusion approach for targeted genome insertion, the DBPs used thus far are still challenging to reprogram, with significant off-target effects. Another approach to alter the target specificity of random transposons is to engineer the transposon DNA sequence. IS608, which is directed by base-pairing interactions between a transposon end and target DNA to insert 3’ to a tetranucleotide sequence, was shown to be targeted more specifically by increasing the length of the guiding sequence contained in the transposon end^29^. However, altering transposon flanking end sequences affects the physical structure and biochemical activity of the transposon, thus limiting the range of viable sequence alterations that can be made.

More recently, protein fusions using a catalytically inactive Cas9 nuclease (dCas9) as an RNA-guided DNA-binding protein have enabled manipulation of genomic DNA and gene expression near user-defined loci^30^. CRISPR interference (CRISPRi) approaches employ a dCas9 protein that uses a synthetic guide RNA (gRNA) to inactivate expression of a particular gene by blocking its transcription through steric hindrance of RNA polymerase^31^. CRISPR activation (CRISPRa) methods use a dCas9-transcription activator fusion that enhances gene expression when targeted to a promoter region^32^. FokI-dCas9 is a nuclease fusion strategy where the FokI nuclease only functions as a dimer and requires targeting of two separate FokI-dCas9 proteins, thus providing increased site-specificity over the Cas9 nuclease^33, 34^. Base editors, which are fusions of a deaminase and dCas9, enable single-base substitutions at gRNA-targeted loci^35,36^. To increase recombination specificity, dCas9 has been used as a fusion to the Gin serine recombinase to better target a particular recombinase recognition site in the human genome^37^. To date however, no dCas9-transposase fusions have been demonstrated to target transposition into a genomic locus; a recent study by Luo et al. showed that a dCas9-piggyBac transposase fusion protein did not have targeted activity^27^.

In this study, we developed a novel system, Cas-Transposon (CasTn) that unites the DNA integration capability of the Himar1 transposase and the programmable genome targeting capability of dCas9 to enable site-specific transposon insertions at user-defined genetic loci. This gRNA-targeted Himar1-dCas9 fusion protein integrates a transposon carrying synthetic DNA payloads into a specific genomic locus (**Figure 1a**), which we demonstrated in both cell-free *in vitro* reactions and in *E. coli*. Cas-Transposon can potentially function in prokaryotic or eukaryotic cells, because the Himar1-dCas9 protein requires no host factors to function. This approach has various uses in metabolic engineering^38^ and emergent gene drive applications^39^.

**Figure 1.**
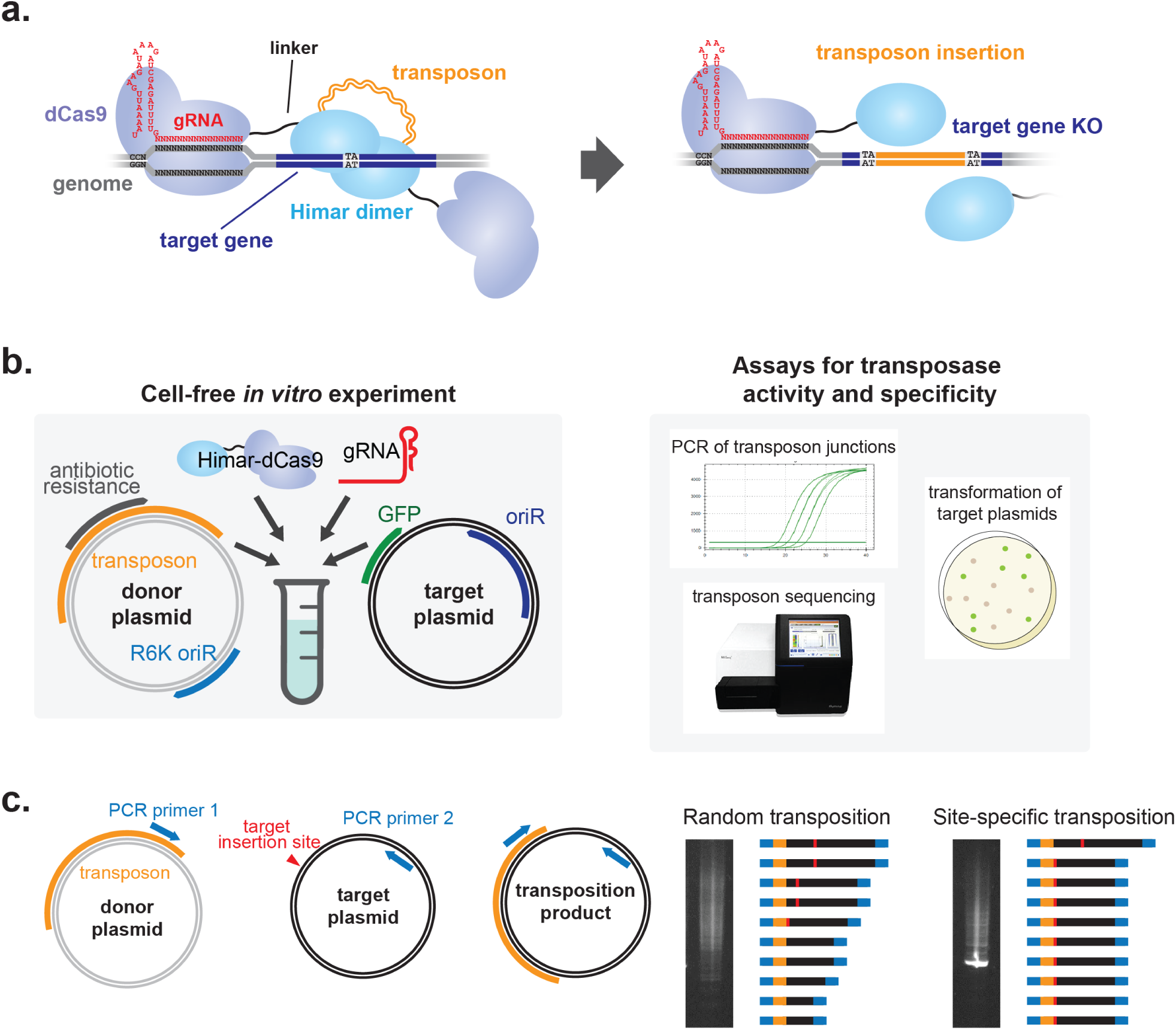
Schematics of *in vitro* Cas-Transposon test system. **(a)** Overview of Himar1-dCas9 protein function. The Himar1-dCas9 fusion protein is guided to the target insertion site by a gRNA, where it is tethered by the dCas9 domain. The Himar1 domain dimerizes with that of another fusion protein to cut-and-paste a Himar1 transposon into the target gene, which is knocked out in the same step. **(b)** Implementation of Cas-Transposon system *in vitro*. Transposon donor and target plasmids were mixed with purified protein and gRNA. Post-transposition target plasmids were analyzed by PCR for plasmid/transposon junctions, transformation and colony analysis, and transposon sequencing. **(c)** Schematic of target plasmid/transposon junction PCR. The PCR was performed using primer 1, which binds the transposon, and primer 2, which binds the target plasmid. Site-specific transposition results in an enrichment for a PCR product corresponding with the expected transposition product.

## MATERIALS AND METHODS

### Strains, media, and growth conditions

All *E. coli* strains were grown aerobically in LB Lennox broth at 37 °C with shaking, with antibiotics added at the following concentrations: carbenicillin (carb) 50 μg/mL, kanamycin (kan) 50 μg/mL, chloramphenicol (chlor) 20-34 μg/mL, and spectinomycin (spec) 240 μg/mL for S17 derivative strains and 60 μg/mL for non-S17 derivative strains. Supplements were added at the following concentrations: diaminopimelic acid (DAP) 50 μM, anhydrotetracycline (aTc) 1-100 ng/mL, and magnesium chloride (MgCl_2_) 20 mM.

### Buffer compositions

Buffers used in the study as follows. Protein Resuspension Buffer (PRB): 20 mM Tris-HCl pH 8.0, 10 mM imidazole, 300 mM NaCl, 10% v/v glycerol. 1 tablet of cOmplete™, Mini, EDTA-free Protease Inhibitor Cocktail (Roche) was dissolved in 10 mL of buffer immediately before use. Protein Wash Buffer (PWB): 20 mM Tris-HCl pH 8.0, 30 mM imidazole, 500 mM NaCl, 10% v/v glycerol. Protein Elution Buffer (PEB): 20 mM Tris-HCl pH 8.0, 500 mM imidazole, 500 mM NaCl, 10% v/v glycerol. Dialysis Buffer 1 (DB1): 25 mM Tris-HCl pH 7.6, 200 mM KCl, 10 mM MgCl_2_, 2 mM DTT, 10% v/v glycerol. Dialysis Buffer 2 (DB2): 25 mM Tris-HCl pH 7.6, 200 mM KCl, 10 mM MgCl_2_, 0.5 mM DTT, 10% v/v glycerol. 10x Annealing Buffer: 100 mM Tris-HCl pH 8.0, 1M NaCl, 10 mM EDTA (pH 8.1).

### Design and construction of the Himar-dCas9 transposase

The design principles of a programmable, site-specific transposition system leverages key insights from previous studies on Himar1 transposases and dCas9 fusion variants^7, 20, 28, 31-36^. The dCas9 protein is a catalytically inactive Cas9 nuclease from *Streptococcus pyogenes* that contains the D10A and H841A amino acid substitutions in its DNA cleavage catalysis sites^7, 31^. This variant has been well-characterized and used as an RNA-guided DNA-binding protein in a number of settings for DNA base editing and transcriptional modulation^31, 32, 35, 36^. Himar1C9 is a hyperactive Himar1 transposase variant that efficiently catalyzes transposition in diverse species and *in vitro*^20^, highlighting its robust ability to integrate without host factors in a variety of cellular environments. The Himar1 transposon features a MmeI restriction site in its inverted repeat sequences, which enables isolation of transposon-insertion site junctions by restriction digest, for facile generation of deep sequencing libraries to interrogate transposon insertion sites. Since the Himar1 transposase only inserts transposons into TA dinucleotide sites, a targeted Himar1-dCas9 fusion protein could potentially enable single-nucleotide precision in insertions, whereas transposases with less well-defined insertion sites (such as Tn5^40^) may not achieve that level of precision targeting. The C-terminus of Himar1C9 is fused to the N-terminus of dCas9 using flexible protein linker XTEN (N-SGSETPGTSESATPES-C), the most optimal linker for FokI-dCas9^33^, because N-terminal dCas9 fusions and C-terminal *mariner*-family transposase fusions have been previously described^28, 33, 34^.

The gene encoding Himar1C9-XTEN-dCas9 (Himar-dCas9) was constructed from the Himar1C9 transposase gene on plasmid pSAM-BT^21^ and the dCas9 gene sequence from pdCas9-bacteria (Addgene plasmid #44249). Linker sequence XTEN was obtained from previously described FokI-dCas9 fusions^33^ and its DNA coding sequence was synthesized as a gBlock® (Integrated DNA Technologies). DNA sequences were PCR amplified using Kapa Hifi Master Mix (Kapa Biosystems) and cloned into expression vectors using NEBuilder® HiFi DNA Assembly Master Mix (New England Biolabs). The Himar-dCas9 gene was cloned into a C-terminal 6xHis-tagged T7 expression vector (yielding plasmid pET-Himar-dCas9), and the protein was purified from Rosetta2 *E. coli* by nickel affinity chromatography to test its activity *in vitro*. Himar-dCas9, dCas9, and Himar1C9 genes were cloned into tet-inducible bacterial expression vectors (yielding plasmids pHdCas9, pdCas9-carb, and pHimar1C9 respectively) to assess protein function *in vivo*. Several tet-inducible bacterial expression vectors for Himar-dCas9 that additionally featured constitutive gRNA expression cassettes were also constructed to evaluate site-specificity of Himar-dCas9 *in vivo*: pHdCas9-gRNA1, pHdCas9-gRNA4, pHdCas9-gRNA5, pHdCas9-gRNA5-gRNA16 containing gRNA_1, gRNA_4, gRNA_5, and gRNA_5 and gRNA_16, respectively. Plasmids used in this study are described in **Table 1**. All gRNAs used in this study are described in **Table 2**.

### Measurement of Himar-dCas9 gene expression knockdown in *E. coli*

We measured expression knockdown of a mCherry reporter gene in *E. coli* strain EcSC83 (MG1655 *galK::mCherry-specR*). Tet-inducible expression vectors pHdCas9-gRNA5-gRNA16 and pdCas9-gRNA5-gRNA16 were used to produce either Himar-dCas9 or dCas9 (a positive control) in each strain along with 2 gRNAs targeting the mCherry gene.

We measured expression knockdown of a GFP reporter gene encoded on the pTarget plasmid in the *E. coli* S17 strain. Tet-inducible expression vectors (pHdCas9-gRNA1, pHdCas9-gRNA4, pHdCas9-gRNA5, pHdCas9 for negative control) were used to express Himar-dCas9 along with a GFP-targeting gRNA in S17 with pTarget.

Saturated overnight *E. coli* cultures were diluted 1:40 into fresh LB media containing aTc (1-100 ng/mL) to induce Himar-dCas9 or dCas9 expression. The induced culture was grown in 200 μL reactions in 96-well plates at 37°C with shaking on a BioTek plate reader. Measurements of OD600 and mCherry (excitation 580 nm, emission 610 nm) and GFP (excitation 485 nm, emission 528 nm) fluorescence levels were taken after 12 hours post-induction.

### Measurement of Himar-dCas9 transposase activity in *E. coli*

Himar-dCas9 and Himar1C9 proteins were expressed in MG1655 *E. coli* from tet-inducible expression vectors pHdCas9 and pHimar1C9, respectively. These strains were conjugated with diaminopimelic acid (DAP) auxotrophic donor strain EcGT2 (S17 *asd::mCherry-specR*)^41^ containing transposon donor plasmid pHimar6, which has a 1.4 kb Himar1 transposon with a chlor resistance cassette and the R6K origin of replication, which does not replicate in MG1655.

Donor and recipient cultures were grown overnight at 37 °C; donors were grown in LB media with DAP and kan, and recipients were grown in LB with carb. 100 uL of donor culture was diluted into 4 mL fresh media and grown for 5 hours at 37 °C. 100 uL of recipient culture was diluted into 4 mL fresh media with 1 ng/mL aTc to induce transposase expression and grown for 5 hours at 37 °C. Donor and recipient cultures were centrifuged and resuspended twice in phosphate-buffered saline (PBS) to wash the cells and quantified on a Nanodrop. 10^9^ donor and 10^9^ recipient cells were mixed, pelleted, resuspended in 20 μL PBS and dropped onto LB agar with 1 ng/mL aTc. The mixed cell droplets were dried at room temperature, and the conjugations were incubated for 2 hours at 37 °C. After conjugation, cells were scraped off, resuspended in PBS by pipetting, and plated on nonselective LB agar plates and LB + chlor (20 μg/mL) plates to select for recipient cells with an integrated transposon. Transposition rates were measured as the ratio of chlor-resistant colony-forming units (CFUs) to total CFUs.

### Purification of Himar-dCas9 protein for *in vitro* transposition assays

His-tagged Himar-dCas9 was purified by nickel affinity chromatography. Plasmid pET-Himar-dCas9 was electroporated into competent Rosetta2 cells (Novagen); transformants were selected on chlor (34 ug/mL) and carb. A single colony was inoculated in 4 mL of LB with chlor (34 ug/mL) and carb and grown to saturation overnight at 37 °C with shaking. 1 mL of saturated culture was diluted into 100 mL of fresh media and grown to OD 0.6-0.8 at 37 °C with shaking. 0.2 mM of Isopropyl β-D-1-thiogalactopyranoside (IPTG) was added to induce Himar-dCas9 protein expression, and the flask was incubated overnight (16 to 18 hours) at 18 °C with shaking. The induced culture was chilled on ice and centrifuged at 4 °C, 7,197 × g for 5 minutes to pellet the cells. The cell pellet was resuspended in 5 mL of Protein Resuspension Buffer (PRB) in a 50 mL conical tube kept chilled in ice water. Cells were lysed using a Qsonica sonicator at 40% power for a total of 120 seconds in 20 second/20 second on/off intervals. The cell suspension was mixed by pipetting, and the sonication cycle was repeated to ensure complete lysis. The lysate was centrifuged at 7,197 × g for 10 minutes at 4 °C to pellet cell debris, and the cleared cell lysate was collected.

To equilibrate the nickel affinity resin, 1 mL of Ni-NTA Agarose (Qiagen) was added to a 15 mL polypropylene gravity flow column (Qiagen), storage buffer was allowed to flow through by gravity, and the resin was washed in 5 mL of PRB and drained. Cleared cell lysate was added to the column and incubated at 4 °C on a rotating platform for 30 minutes. The cell lysate was flowed through, and the nickel resin was washed with 25 mL PWB in 5 mL increments. The protein was eluted with PEB in 5 fractions of 0.5 mL each. Each elution fraction was analyzed by running an SDS-PAGE gel; most protein eluted in fractions 2-4. Elution fractions 2-4 were combined and dialyzed overnight in 500 mL of DB1 at 4 °C using 10K MWCO Slide-A-Lyzer™ Dialysis Cassettes (Thermo Fisher). The protein was dialyzed again in 500 mL DB2 for 6 hours. The dialyzed protein was quantified with the Qubit Protein Assay Kit (Thermo Fisher) and divided into single-use aliquots that were snap frozen in dry ice and ethanol and stored at −80 °C.

### *In vitro* transposition reaction using purified Himar-dCas9 protein

We characterized the specificity and efficiency of transposition by Himar-dCas9 within *in vitro* reactions (**Figure 1b**). Each *in vitro* Himar-dCas9 transposition reaction was performed in a buffer consisting of 10% glycerol, 2 mM dithiothreitol (DTT), 250 μg/mL of bovine serum albumin (BSA), 25 mM HEPES (pH 7.9), 100 mM NaCl, and 10 mM MgCl_2_. *In vitro* transposition reactions contained up to 5 nM each of transposon donor plasmid pHimar6 and target plasmid pGT-B1, up to 100 nM of transposase/gRNA complexes, and up to 800 ng of background *E. coli* genomic DNA in a final volume of 20 μL. pGT-B1 is a 6 kb minimal plasmid containing the pBBR1 origin of replication, a beta-lactamase selection marker, and a constitutively expressed *GFP* gene^41^.

Plasmid DNA was purified using the ZymoPureII midiprep kit (Zymo Research). *E. coli* genomic DNA was purified using the MasterPure Gram Positive DNA Purification Kit (Epicentre). All DNAs were purified again using the Zymo Clean and Concentrator-25 (Zymo Research) kit to remove all traces of RNAse and quantified using the Qubit dsDNA BR Assay kit (Invitrogen). The gRNAs were synthesized *in vitro* using the GeneArt™ Precision gRNA Synthesis Kit (Invitrogen) according to manufacturer’s instructions and were aliquoted into single-use tubes, flash frozen in dry ice and ethanol, and stored at −80 °C until usage. gRNA concentrations were measured using the Qubit RNA HS Assay Kit (Invitrogen).

To set up *in vitro* reactions, frozen aliquots of transposase protein and gRNAs were thawed on ice. The protein was diluted to 20x final concentration in DB2 buffer, and gRNAs were diluted to the same molarity in nuclease-free water. The diluted protein and gRNA were mixed in equal volumes and incubated at room temperature for 15 minutes. Transposon donor DNA, target plasmid DNA, and background DNA (if applicable) were mixed on ice with 10 μL of a 2× buffer master mix and water to reach a volume of 18 μL. 2 μL of the protein/gRNA mixture was added last to the reaction. In reactions where the transposase/gRNA complex was preloaded onto the target plasmid, the target plasmid was mixed with protein and gRNA and incubated at 30 °C for 10 minutes, and the donor DNA was added last. Transposition reactions were incubated for 3 to 72 hours at 30-37 °C and then heat-inactivated at 75 °C for 20 minutes. Transposition products were purified using magnetic beads^42^, eluted in 45 μL of nuclease-free water, and stored at −20 °C.

### qPCR assay for site-specific insertions

One method used to evaluate the specificity and efficiency of Himar-dCas9 within *in vitro* transposition reactions was a series of qPCRs (**Figure 1b, c**). For each reaction, 2 qPCRs were performed to obtain the measure of *relative Cq*: one PCR amplifying transposon-target plasmid junctions, and another PCR amplifying the target plasmid backbone to normalize for template DNA input across samples (**Figure 1c**). Relative Cq measurements shown in this study are the differences between these two Cq values. For in *vitro* transposition into pGT-B1 (target plasmid used in *in vitro* experiments), primers p433 and p415 were used for junction PCRs, and primers p828 and p829 were used for control PCRs. For *in vitro* transposition into pTarget (target plasmid also used for *in vivo* bacteria experiments), primers p898 and p415 were used for junction PCRs and primers p899 and p900 were used for control PCRs. All qPCR primers used in this study are listed in **Table 3**.

PCR reactions contained 1 μL each of 10 μM forward and reverse primers, 1 μL of purified transposition products as template DNA, 7 μL water, and 10 μL of Q5 2X Master Mix (NEB) + SYBR Green. Reactions were thermocycled using a Bio-Rad C1000 touch qPCR machine for 1 minute at 98 °C, followed by 35 cycles of 98 °C denaturation for 10 seconds, 68 °C annealing for 15 seconds, and 72 °C extension for 2 minutes. PCR products were quantified by SYBR Green fluorescence and qualitatively analyzed for specificity on agarose DNA gels.

### Transformation assay for *in vitro* transposition reaction products

A second method used to measure transposition specificity and efficiency was transformation of the reaction product DNA into competent *E. coli* and analyzing transposon inserts in individual transformants (**Figure 1b**). 5 μL purified DNA from an *in vitro* transposition reaction was mixed with 45 μL distilled water and chilled on ice. 10 μL of thawed MegaX electrocompetent *E. coli* (Invitrogen) was added and mixed by pipetting gently. The mixture was transferred to a 0.1 cm gap electroporation cuvette (BioRad) and electroporated at 1.8 kV. Cells were recovered in 1 mL SOC and incubated with shaking at 37 °C for 90 minutes. The cells were plated on antibiotic selection plates with chlor (34 ug/mL) to select for target plasmids (pGT-B1) containing transposons, and on plates with carb to measure the electroporation efficiency of pGT-B1. The efficiency of transposition was measured as the ratio of chlor-resistant transformants to carb-resistant transformants. To assess specificity of inserted transposons, we performed colony PCR on transformants using the primer set p433/p415 with KAPA2G Robust HotStart ReadyMix (Kapa Biosystems) to amplify junctions between the Himar1 transposon from pHimar6 and the pGT-B1 target plasmid, which were analyzed by Sanger sequencing.

### Transposon sequencing library preparation

To quantitatively survey the distribution of transposition events performed by Himar-dCas9 in *in vitro* reactions, we performed transposon sequencing on reaction products (**Figure 1b, Suppl. Figure S2**). Transposon junctions were PCR amplified from *in vitro* transposition reactions using primer sets p923/p433 and p923/p922, using Q5 HiFi 2x Master Mix (NEB) + SYBR Green. Primer p923 binds the Himar1 transposon from pHimar6, while p433 and p922 bind to target plasmid pGT-B1. Reactions were thermocycled using a Bio-Rad C1000 touch qPCR machine for 1 minute at 98 °C, followed by cycles of 98 °C denaturation for 10 seconds, 68 °C annealing for 15 seconds, and 72 °C extension for 2 minutes. PCR reactions were stopped in late exponential phase in order to avoid oversaturation of PCR products. PCR products were purified using magnetic beads^42^, and 100-200 ng of DNA per sample was digested with MmeI (NEB) for 1 hour in a reaction volume of 40 μL. The MmeI digestion products were purified using Dynabeads M-270 Streptavidin beads (ThermoFisher) according to manufacturer instructions. The digested transposon ends, bound to magnetic Dynabeads, were mixed with 1 μg sequencing adapter DNA (see next section), 1 μL T4 DNA ligase, and T4 DNA ligase buffer in a total reaction volume of 50 μL. The ligation reactions were incubated at room temperature (∼23 °C) for 1 hour; the ligase enzyme and excess sequencing adapter were removed by washing the beads according to Dynabead manufacturer instructions. Dynabeads were resuspended in 40 μL of water.

2 μL of the Dynabeads were used as template for the final PCR using barcoded P5 and P7 primers and Q5 HiFi 2× Master Mix (NEB) + SYBR Green. Reactions were thermocycled using a Bio-Rad C1000 touch qPCR machine for 1 minute at 98 °C, followed by cycles of 98 °C denaturation for 10 seconds, 67C annealing for 15 seconds, and 72 °C extension for 20 seconds. PCR reactions were stopped in late exponential phase in order to avoid oversaturation of PCR products. Equal amounts of DNA from all PCR reactions were combined into one sequencing library, which was purified and size-selected for 145 bp products using the Select-a-Size Clean and Concentrator kit (Zymo). The library was quantified with the Qubit dsDNA HS Assay Kit (Invitrogen) and combined in a 7:3 ratio with PhiX sequencing control DNA. The library was sequenced using a MiSeq V2 50 Cycle kit (Illumina) with custom read 1 and index 1 primers spiked into the standard read 1 and index 1 wells. Reads were mapped to the pGT-B1 plasmid using Bowtie 2^43^.

### Construction of sequencing adapter by annealing

Oligonucleotides Adapter_T and Adapter_B were diluted to 100 μM in nuclease-free water. 10 μL of each oligo was mixed with 2.5 μL water and 2.5 μL of 10× Annealing Buffer. Using a thermocycler, the mixture was heated to 95 °C and slowly cooled at 0.1C/second to 4 °C to yield 25 μL of 40 μM sequencing adapter. The adapter was stored at −20 °C until ready for use.

### *In vivo* assays for transposition into a target plasmid

S17 *E. coli* were sequentially electroporated with plasmid pTarget as a target plasmid and then one of several pHdCas9-gRNA plasmids (pHdCas9-gRNA1, pHdCas9-gRNA4, pHdCas9-gRNA5, or pHdCas9), which are bacterial expression vectors for Himar-dCas9 and a gRNA (**Figure 4a, Table 1**).Transformants were selected on LB with carb and spec (240 μg/mL). Transformants were grown from a single colony to mid-log phase in liquid selective media, electroporated with 130 ng pHimar6 transposon donor plasmid DNA, and recovered in 1 mL LB for 1 hour at 37C with shaking post-electroporation. 100 μL of a 10^−3^ dilution of the transformation was plated on LB agar plates with spec (240 μg/mL), carb, chlor (20 ug/mL), MgCl_2_ (20 mM), and aTc (0-2 ng/mL). Plates were grown at 37 °C for 16 hours. Between 10^3^ and 10^4^ colonies were scraped off each plate into 2 mL PBS and homogenized by pipetting. 500 uL of the cells were miniprepped using the QIAprep kit (Qiagen).

Minipreps from each transformation were evaluated by qPCR for junctions between the transposon from pHimar6 and the pTarget plasmid and by a transformation assay. qPCR assays for transposon-target plasmid junctions were performed as described above, using primers p898 and p415 and 10 ng of miniprep DNA as PCR template. The control PCR to normalize for pTarget DNA input was performed with primers p899 and p900. In transformations, 150 ng of plasmid DNA was electroporated into 10 uL of MegaX electrocompetent cells diluted in 50 uL of ice-cold distilled water. Cells were immediately recovered in 1 mL of LB and incubated with shaking at 37 °C for 90 minutes. The cells were plated on LB agar with chlor (20 ug/mL) and spec (60 ug/mL) to select for pTarget plasmids containing a transposon from pHimar6. We performed colony PCR using the primer set p898/p415 with KAPA2G Robust HotStart ReadyMix (Kapa Biosystems) to amplify transposon-pTarget junctions, which were analyzed by Sanger sequencing.

## RESULTS

### Establishing DNA-targeting and transposition activity of Himar-dCas9

Because Himar1C9-dCas9 is a novel synthetic fusion protein, we first verified that both the Himar1 and dCas9 components remained functional when fused. To check that Himar-dCas9 was still capable of binding a DNA target specified by a gRNA, we expressed Himar-dCas9 in an *E. coli* strain with a genomically integrated mCherry gene, along with 2 gRNAs targeting mCherry (gRNA_5 and gRNA_16 in **Table 2**). If Himar-dCas9 bound to the mCherry gene, the protein would sterically hinder RNA polymerase and decrease transcription, resulting in decreased mCherry fluorescence. Indeed, we observed mCherry knockdown when Himar-dCas9 was expressed, indicating that the fusion protein’s DNA binding functionality was intact (**Suppl. Figure S1a**). To verify Himar-dCas9 transposase activity, we conjugated a Himar1 transposon with a chloramphenicol resistance gene (on plasmid pHimar6) from EcGT2 donor *E. coli* into MG1655 *E. coli* expressing Himar-dCas9 (but no gRNA) from plasmid pHdCas9, or Himar1C9 transposase from plasmid pHimar1C9. We measured the rate of transposition as the proportion of cells that acquired a genomically integrated transposon (**Suppl. Figure S1b**). We confirmed that Himar-dCas9 mediates transposition events in *E. coli*, although at a lower rate (about 2 log-fold) compared with Himar1C9, which may be associated with lower expression of Himar-dCas9, which is a much larger and metabolically costly protein to produce, or with altered DNA affinity by dCas9 even in the absence of gRNA^44^.

### An *in vitro* reporter system to assess targeted transpositions by Himar-dCas9

To best establish and optimize the parameters for efficient targeted transposition reactions, we next developed an *in vitro* reporter system to explore the targeted transposition activity of Himar-dCas9. Purified Himar-dCas9 protein was mixed with a transposon donor pHimar6 plasmid (containing a Himar1 transposon with a chlor-resistance gene), a transposon target pGT-B1 plasmid (containing a *GFP* gene), and one or more gRNAs targeted to various loci along *GFP* (**Figure 1b, Tables 1-2**). We analyzed transposon insertion events into the pGT-B1 plasmid by several assays. First, quantitative PCR (qPCR) of target plasmid/transposon junctions, using one primer designed to anneal to a part of the transposon DNA and one primer designed to anneal to a part of pGT-B1, enabled quantification of transposons that were inserted into the target plasmid, as well as qualitative assessment of transposition specificity based on enrichment of qPCR products of the expected amplicon size (**Figure 1b-c, Table 3**). For every transposon/target junction qPCR, we also performed a control qPCR that amplifies the pGT-B1 plasmid’s backbone to control for variations in DNA input between samples. Relative Cq measurements were taken as the difference between the Cq values from the junction and control qPCR reactions. Next-generation transposon sequencing (Tn-seq) further enabled quantitative measurement of the distribution of inserted transposons within the target plasmid (**Suppl. Figure S2**). Finally, transposition reaction products were transformed into competent *E. coli* to further probe specificity of transposition insertion sites. Because the donor pHimar6 plasmid has a R6K origin of replication that is unable to replicate in *E. coli* without the *pir* replication gene, we phenotypically selected for transformants containing the target pGT-B1 plasmid with an integrated transposon. Transposition efficiency was determined by dividing the number of chlor-resistant transformants (CFUs with a target plasmid carrying a transposon) by the number of carbenicillin-resistant transformants (total CFUs with a target plasmid). Sanger sequencing of the target plasmid from chloramphenicol-resistant transformants revealed the site of integration and the transposition specificity.

### Efficiency and site-specificity of Himar-dCas9 transposon insertions is gRNA-dependent

Using the *in vitro* reporter system, we first assessed how the genomic orientation and distance of the gRNA relative to the target TA dinucleotide site affects site-specificity of transposition. We designed gRNAs spaced 5 to 18 bp from a target TA site, targeting either the template strand or non-template strand of the GFP gene (**Figure 2a, Table 1**). Each individual gRNA design was tested in triplicate *in vitro* reactions using 30 nM of purified Himar-dCas9 protein, 30 nM of gRNA, 2.27 nM of pHimar6, and 2.27 nM of pGT-B1.

**Figure 2.**
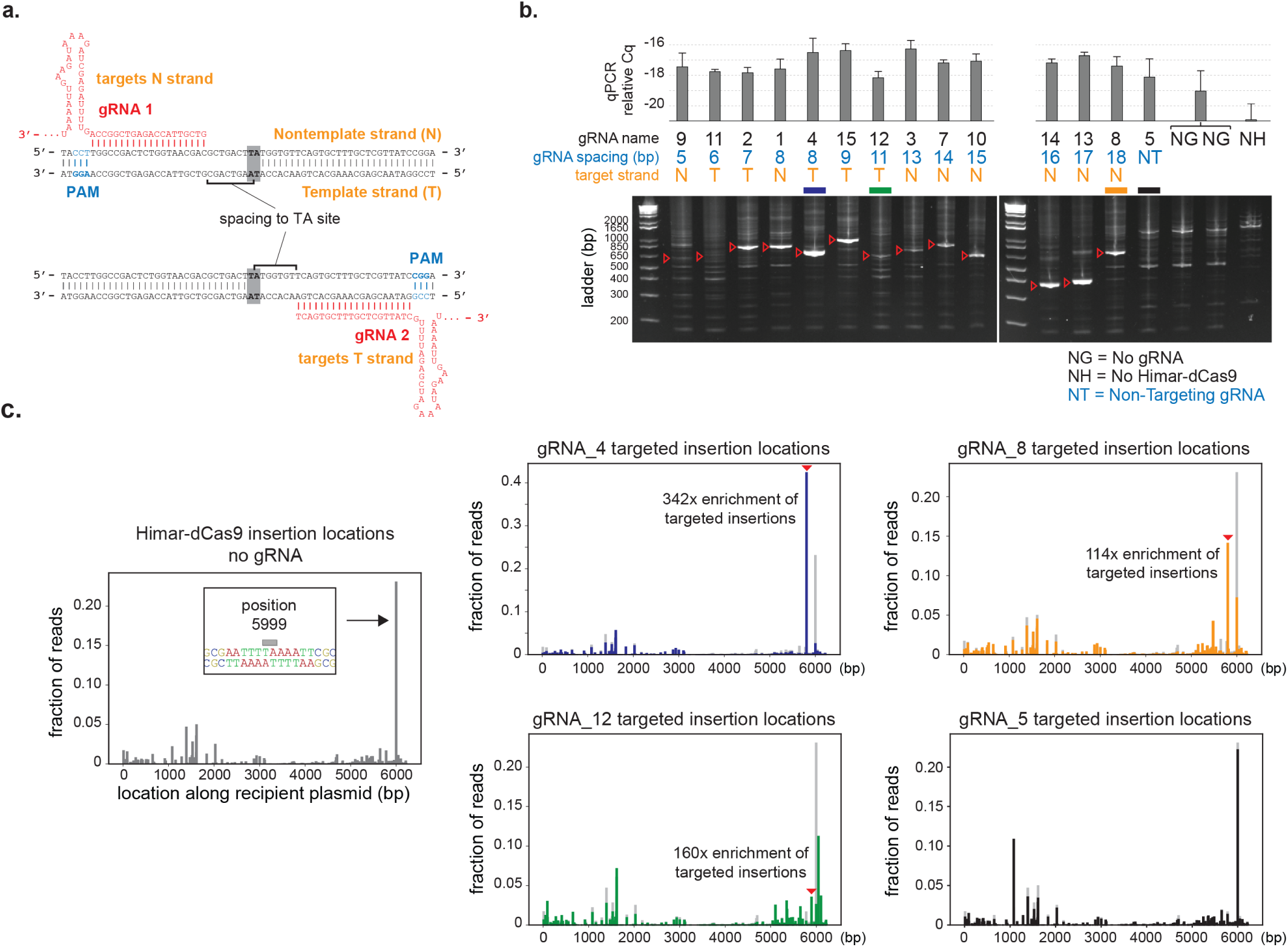
Himar-dCas9 specificity is dependent on gRNA spacing and target site. **(a)** Illustration of gRNA strand orientation and spacings to TA insertion site. **(b)** PCR analysis of transposon/target junctions from *in vitro* reactions containing 30 nM Himar-dCas9/gRNA complex, 2.27 nM transposon donor DNA, and 2.27 nM target DNA. Reactions (n=3) were run using gRNAs with spacings between 5 and 18 bp from the TA insertion site. Non-targeting gRNA (gRNA5), no gRNA, and no transposase controls were also performed. Red arrows indicate expected site-specific PCR products for each gRNA. Error bars indicate standard deviation. **(c)** Transposon sequencing results for reactions with no gRNAs (left, gray, n=4) or with gRNA_4 (blue, n=3), gRNA_8 (orange, n=3), gRNA_12 (green, n=3), or gRNA_5 (black, n=3). The baseline random distribution of transposons along the recipient plasmid in each panel with a gRNA is shown in light gray.

Using the qPCR assay, we found that a single gRNA is sufficient to mediate site-specific transposition (**Figure 2b**). Furthermore, the site-specificity of transposition appears to be dependent on the gRNA spacing to the target TA site. All gRNA-targeted insertion events occurred at the nearest TA distal to the 5’ end of the gRNA, independent of the targeted DNA strand (**Figure 2b**). We found a significant preference for site-specific transpositions for gRNAs with 7-9 bp and 16-18 bp spacings, based on the observed strong PCR bands of expected amplicon size for on-target transpositions. At very short spacings (5-6 bp), there was no PCR band of the expected amplicon junction length, suggesting that the Himar-dCas9 protein sterically hinders transposition at distances less than 7 bp from the target TA site. At spacings between 11-15 bp, there was a faint expected PCR band, indicating that transposition at those sites may occur, but site-specific transposition was not optimal. These findings are consistent with the previously observed spacing dependence for FokI-dCas9 proteins built using the same XTEN peptide linker^33^. The bimodal distribution of robustly targeting gRNA spacings may be due to the DNA double helix providing steric hindrance at intermediate spacings, since the spacing peaks are approximate 1 helix turn (∼10 bp) apart. Comparison of relative Cq values between gRNA-bound Himar-dCas9 versus Himar-dCas9without a gRNA suggests that transposition events occur at a higher rate in the presence of a targeting gRNA (**Figure 2b**). This observation may be the result of gRNA-targeted Himar-dCas9 transposases binding near potential TA insertion sites, thus circumventing the need for a transposase dimer loaded with a transposon to search for TA sites.

To assess the distribution of transposon insertions around the target pGT-B1 plasmid, we performed transposon sequencing on transposition products resulting from 3 GFP-targeting gRNAs (gRNA_4, gRNA_8 and gRNA_12), a non-targeting gRNA (gRNA_5), and no gRNA (**Figure 2c**). The baseline distribution of random transposon insertion events was generated from quadruplicate *in vitro* reactions with no gRNA added. Random transposon insertions were present throughout the 6.2 kb pGT-B1 plasmid, with a spike in transposition abundance at position 5999, which is a TA site in the middle of a 12 bp stretch of T/A nucleotides. This result is consistent with the observation that Himar1 transposase preferentially inserts transposons into flexible, T/A-rich DNA^45^. In contrast, gRNA-targeted insertions were less likely to be inserted into position 5999 and were enriched at their respective gRNA-adjacent TA sites, compared with baseline, to varying degrees (**Figure 2b-c**). The best targeted insertion was found using gRNA_4, with an optimal spacing of 8 bp from the target TA site, had 42% of all transposon insertions being exactly at the target site, a 342-fold enrichment over baseline. Comparison of targeted insertion fold-enrichment across different gRNAs suggests that the specific target site and flanking DNA play a role in the specificity of transposon integration. For instance, gRNA_12 had a higher fold-enrichment of insertions at its target site than gRNA_8, but a lower fraction of insertions, suggesting that the lack of specificity of gRNA_12 may be attributable to its target site being disfavored for transposition. Together, these results further show that Himar-dCas9 mediates targeted transposon insertion to an intended integration site with the help of an optimally spaced gRNA.

Given that *mariner* transposases dimerize in solution in the absence of DNA^46^, we hypothesized that Himar-dCas9 dimerizes spontaneously, and then the active Himar1 dimer is guided to a gRNA-specific target locus by one of the dCas9 domains in the Himar-dCas9 dimer (**Figure 1a**). This mechanism of activity is consistent with the observation that a single gRNA is sufficient to direct targeted transposition. Further support for this hypothesis comes from *in vitro* reactions containing pairs of gRNAs targeting the same TA site, but on opposite complementing strands (**Suppl. Figure S3**). If Himar1 subunits did not spontaneously dimerize, then dimerization of Himar-dCas9 would be enhanced by loading two Himar-dCas9 monomers onto the same target plasmid, so that pairs of Himar1 monomers would be tethered in close proximity. We set up reactions in which target DNA was first preloaded with either paired or single gRNA/Himar-dCas9 complexes and then mixed with transposon donor DNA (**Suppl. Figure S3a**). In these experiments, the final reaction contained 5 nM of Himar-dCas9 protein, 5 nM of donor DNA, 5 nM of target DNA, and 2.5 nM of each of two gRNAs. We observed no difference in transposition rate or specificity between the gRNA/Himar-dCas9 complexes preloaded as pairs or as singletons (**Suppl. Figure S3b, c**). The observation that preloading pairs of Himar-dCas9 complexes does not improve transposition is consistent with the hypothesis that transposase dimers formed before one of the gRNA/dCas9 domains targeted the dimer to its final location along the DNA.

### Site-specific transposition by Himar-dCas9 is robust across a range of protein and DNA concentrations

To assess the robustness of Himar-dCas9 to various experimental conditions and determine the optimal parameters for site-specific transposition, we explored different concentrations of (1) protein-gRNA complexes, (2) transposon donor plasmid (pHimar6) DNA, (3) target plasmid (pGT-B1) DNA, and (4) background off-target DNA within *in vitro* transposition reactions containing a single gRNA (gRNA_4). We also performed *in vitro* reactions over different temperatures and reaction times.

We first varied concentrations of Himar-dCas9/gRNA complexes and found that site-specific transposition could be detected by our PCR assay in *in vitro* reactions with at least 3 nM of Himar-dCas9/gRNA complexes, using 5 nM of donor and 5 nM of target plasmids (**Figure 3a**). Increasing the Himar-dCas9/gRNA concentration appeared to increase the yield of targeted transposition events. We further purified and transformed the target pGT-B1 plasmid from these reactions into electrocompetent *E. coli* to assess the efficiency and specificity of transposition (**Figure 3b**); the trend of higher transposition at higher transposase concentration is consistent with qPCR measurements. At 30 nM of the Himar-dCas9/gRNA complex, the specificity of transposon insertion into the targeted TA site was 44% (11 of 25 colonies had correct insertion site). The specificity of insertion at 100 nM of the complex did not further increase and remained at 47.5% (19 of 40 colonies with correct insertions).

**Figure 3.**
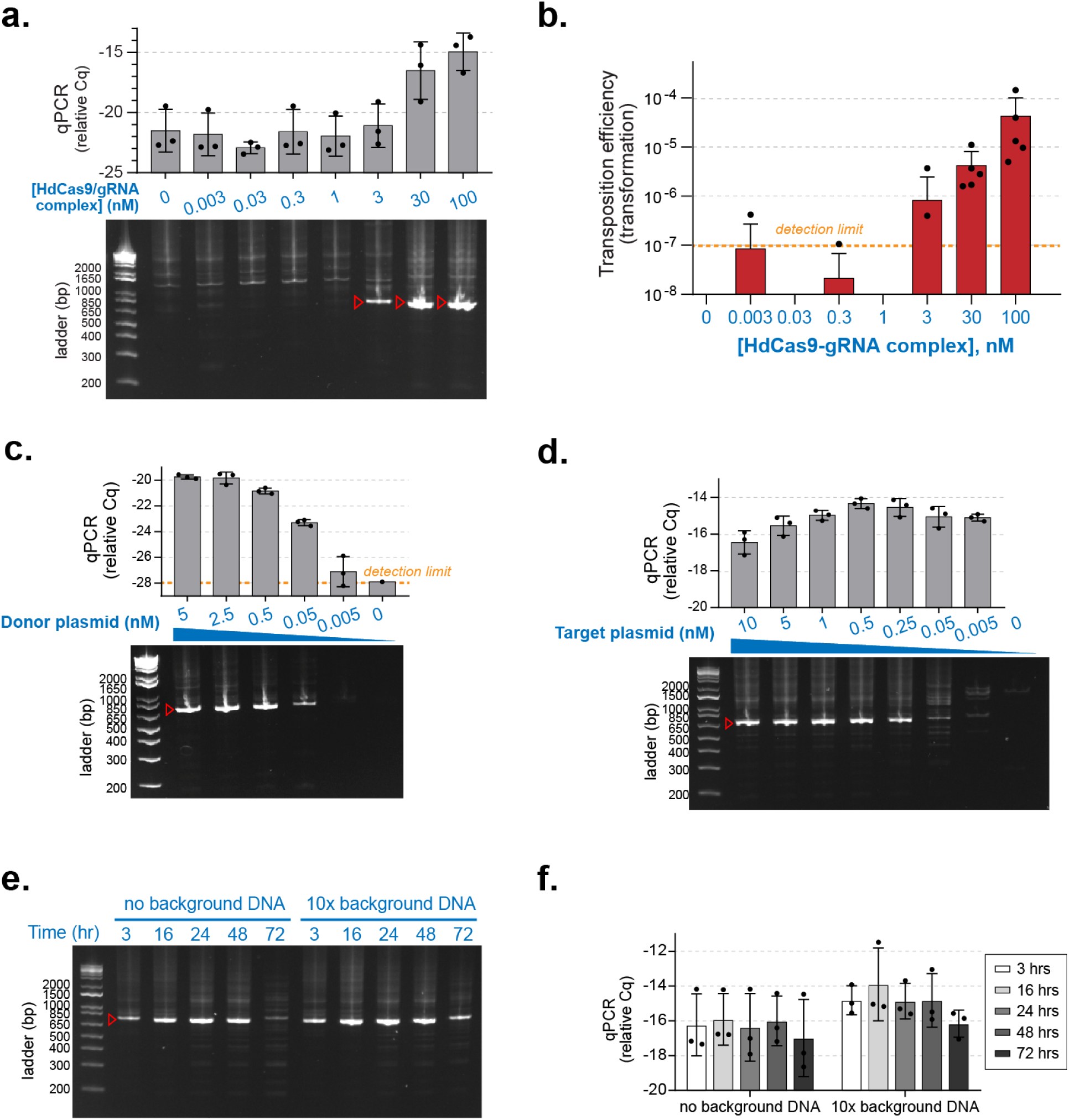
Himar-dCas9-mediated site-specific transposition is robust to changes in ribonucleoprotein complex and DNA concentration. Target plasmids were pGT-B1 and donor plasmids were pHimar6. **(a)** PCR analysis of transposition reactions (n=3) using varying levels of Himar-dCas9/gRNA_4 complexes. Reactions were performed for 3 hours at 30 °C with 5 nM of donor and recipient plasmid DNA. **(b)** Transformation assay to measure transposition rates in reactions using varying levels of Himar-dCas9/gRNA_4 complexes (n=5). Reactions were performed for 3 hours at 30 °C with 5 nM of donor and recipient plasmid DNA.. **(c)** PCR analysis of transposition reactions (n=3) using varying levels of donor plasmid DNA. Reactions were performed for 3 hours at 30 °C with 5 nM of recipient plasmid DNA and 30 nM Himar-dCas9/gRNA_4 complex. **(d)** PCR analysis of transposition reactions (n=3) using varying levels of recipient plasmid DNA. Reactions were performed for 3 hours at 30 °C with 0.5 nM of donor plasmid DNA and 30 nM Himar-dCas9/gRNA_4 complex. **(e)** PCR analysis of transposition reactions (n=3) performed for different lengths of time, in the presence or absence of background non-specific DNA. Reactions were performed at 37 °C with 1 nM of recipient plasmid DNA, 1 nM of donor plasmid DNA, and 100 nM Himar-dCas9/gRNA_4 complex. Background *E. coli* genomic DNA was present at 10×the mass of recipient plasmid DNA. **(f)** qPCR measurement of transposition efficiency in reactions shown in panel (e). n=3 for each reaction condition. In all panels, red arrows indicate the expected PCR product for gRNA_4, and error bars indicate standard deviation. Cq measurements correspond to log-scale differences in transposase activity.

**Figure 4.**
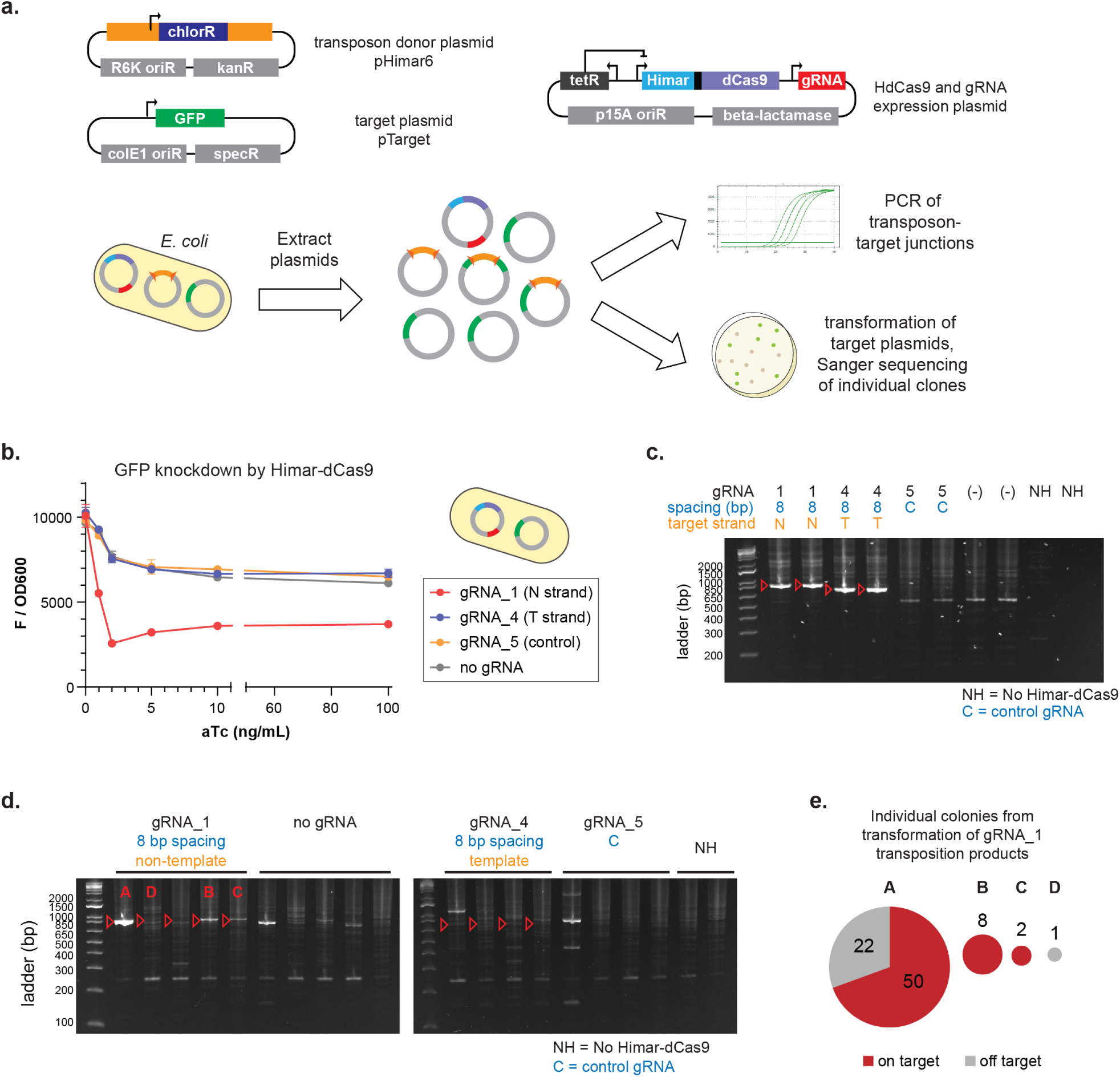
Himar-dCas9 performs site-specific transpositions into plasmids in *E. coli*. **(a)** Three plasmids were transformed into S17 *E. coli* to create a testbed for Himar-dCas9 transposition specificity *in vivo*. Post-transposition plasmids were extracted from the bacteria and analyzed by PCR and by transformation into competent *E. coli* with Sanger sequencing of plasmids from individual colonies. **(b)** Himar-dCas9 knocks down GFP expression from the pTarget plasmid *in vivo* in *E. coli* with gRNA_1, which targets the non-template strand (N) of the GFP gene. Himar-dCas9 does not knock down GFP fluorescence when expressed with a gRNA complementing the template strand (T) or with a non-targeting gRNA (NT) or no gRNA. n=2 per gRNA and ATC concentration; error bars indicate standard deviation. **(c)** PCR assay of *in vitro* transposition reactions using donor plasmid pHimar6 and recipient plasmid pTarget. 2.27 nM of donor and recipient plasmids along with 30 nM Himar-dCas9/gRNA complex were incubated for 3 hours at 30C. Expected PCR products of targeted insertions are shown with red arrows. **(d)** PCR analysis of pTarget-transposon junctions resulting from *in vivo* transposition in bacteria. 3/5 of gRNA_1 PCR products show enrichment for the targeted insertion product. Transpositions A, B, C, and D with gRNA_1 were also analyzed by transformation and colony analysis. **(e)** Plasmid pools from 4 independent *in vivo* transposition experiments using gRNA_1 were transformed into *E. coli*, and the resultant colonies were analyzed by PCR and Sanger sequencing. The pie charts show the number of colonies containing on-target and off-target transposition products from each plasmid pool, with chart area proportional to the total number of colonies.

Next, we explored whether site-specific transposition was affected by DNA concentrations of the donor or target plasmid. Using 5 nM of the target plasmid DNA, the transposition activity was robust across 0.05 nM to 5 nM of donor plasmid DNA, with greater rates of transposition occurring at higher donor DNA concentrations (**Figure 3c**). Similarly, using 0.5 nM of donor plasmid DNA, site-specific transposition was observed across a wide range of target plasmid concentrations from 0.25 nM to 10 nM (**Figure 3d**). While the absolute rate of transposition (as assessed by Cq of the transposon-target junction qPCR) was higher at higher donor DNA concentrations, the relative Cq remained relatively constant across target DNA concentrations, indicating that a similar proportion of target plasmids received a transposon in each reaction.

We also tested whether the gRNA-guided Himar-dCas9 could efficiently transpose into a targeted site in the presence of background DNA and whether the amount of transposition changed over longer reaction times. We added up to 10 times more background *E. coli* genomic DNA than target plasmid DNA to reactions containing 10 nM of Himar-dCas9/gRNA, 1 nM of donor DNA, and 1 nM of target DNA. Across different ratios of target to background DNA concentrations tested, Himar-dCas9 was able to locate the gRNA-targeted site and insert transposons site-specifically with no observed loss of specificity or efficiency (**Suppl. Figure S4a**). When this transposition reaction with background DNA was performed at 37 °C and over longer time courses instead of the standard protocol of 30 °C for 3 hours, to mimic conditions in living cells growing at 37 °C, we observed similar results (**Suppl. Figure S4b-c, Figure 3e-f**). In the presence of 10 times more background DNA than target plasmid DNA, 10 nM and 100 nM of Himar-dCas9-gRNA were capable of mediating site-specific transposition (**Suppl. Figure S4b-c, Figure 3e-f**). The relative Cq and PCR band intensity of transposon-target junctions increased slightly between 3 and 16 hours, but PCR bands remained the same size, suggesting that gRNA-guided transposases are faster at locating the target site than catalyzing transposition and that the increase in site-specific transposon insertions over time is performed by gRNA-dCas9 bound transposases. After 16 hours, site-specific transposition events appeared to reach a plateau; the slight loss of specific transposon-target junctions observed at 72 hours by PCR is likely due to degradation of the *in vitro* reaction components and products over time (**Suppl. Figure S4b, Figure 3e**).

Together, these results highlight that Himar-dCas9/gRNA mediates targeted transposon insertions across a range of experimental conditions for *in vitro* reactions, including physiologically relevant temperatures (30-37 °C) and protein/DNA concentrations. In bacteria, 1 nM corresponds to approximately 1 molecule per cell, while in eukaryotic cells, 1 nM corresponds to approximately 1000 molecules per cell^47^. Targeted transposition was observed to occur at protein concentrations of 1-100 nM (1-100 molecules of protein per bacterial cell) and DNA concentrations of less than 1 to 10 nM (1-10 DNA copies per bacterial cell). In bacteria, these concentrations are physiologically achievable with low protein expression and with transposon donor/target DNA present as a single chromosomal copy or on a low/medium copy number plasmid. Notably, we did not experimentally find an upper limit of protein/DNA concentrations for effective targeted transposition beyond the loss of specific targeting due to increased background transpositions; nevertheless, the CasTn system can be used with different plasmid expression systems to modulate copy numbers of both protein and DNA.

### Himar-dCas9 mediates site-specific transposon insertions into plasmids *in vivo*

Since Himar-dCas9 robustly facilitated site-specific transposon integration *in vitro*, we tested the ability of the system to make site-specific transpositions in *E. coli* bacteria. In the first system, we transformed a set of 3 plasmids into S17 *E. coli*: pTarget, which contains a GFP target gene; pHimar6, the transposon donor plasmid; and a tet-inducible expression vector for Himar-dCas9 that also contained 0 or 1 gRNA targeting GFP (**Figure 4a**). We grew the cells containing these 3 plasmids for 16 hours on selective agar plates with MgCl_2_ and anhydrotetracycline (aTc) to allow for transposition to occur, and then extracted all plasmids from these cells. Transposition specificity in these plasmid pools were determined by two methods: PCR of transposon-target plasmid junctions, and transformation of the pooled plasmids into competent cells and analysis of transposon insertions in individual transformants.

We first verified that the Himar-dCas9 system components functioned *in vivo* in our assay strain. The S17 strain was transformed with pTarget and one of several Himar-dCas9/gRNA expression vectors for gRNA_1, gRNA_4, non-targeting gRNA_5, or no gRNA. By measuring Himar-dCas9-mediated knockdown of GFP in these strains, we confirmed that gRNAs targeted Himar-dCas9 to the pTarget plasmid as expected and determined the optimal concentration of aTc for inducing Himar-dCas9 expression (**Figure 4b**). Consistent with previously reported results^31^, gRNA_1, which targets the non-template strand of the GFP gene, caused knockdown of GFP expression, but gRNA_4, which targets the template strand and does not sterically hinder RNA polymerase, did not cause GFP knockdown. Himar-dCas9 concentrations reached saturation at aTc induction levels of 2 ng/mL, as further increasing the concentration of aTc did not result in further knockdown of GFP by gRNA_1. We also validated that purified Himar-dCas9 protein with gRNA_1 or gRNA_4 mediated targeted transposition into the GFP gene of pTarget *in vitro* (**Figure 4c**).

In the *in vivo* assay, S17 *E. coli* containing pTarget and a Himar-dCas9/gRNA expression vector were transformed with transposon donor plasmid pHimar6. After this transformation, cells were plated on selective agar plates containing a saturating concentration of MgCl_2_ to enable transposase activity and 1 ng/mL aTc to induce expression of Himar-dCas9 while avoiding overproduction inhibition of the Himar1C9 transposase^48^. After 16 hours of growth at 37 °C, we extracted plasmids from the pooled colonies and analyzed the plasmid pools for site-specific transposon insertion events. PCR for transposon-target plasmid junctions showed that gRNA_1 produced detectable site-specific transposon insertions into pTarget in 3 out of 5 independent replicates (**Figure 4d**). gRNA_4, however, did not produce an enrichment of PCR products corresponding to its targeted insertion site.

We further evaluated site-specificity of transposition by transforming the plasmid pools into electrocompetent MegaX *E. coli* and analyzing individual transformants by colony PCR and Sanger sequencing, to confirm that Himar-dCas9 with gRNA_1 mediated precisely targeted transposon insertions into pTarget. Analyzing 4 independent plasmid pools from cells that expressed gRNA_1, we found that 3 out of 4 transformations produced colonies with mostly or all site-specific transposition products (**Figure 4e**). In transformations of 4 plasmid pools from cells without a gRNA, we did not obtain any transformants containing pTarget plasmids with an integrated transposon. Taken together, these results demonstrates *in vivo* targeted transposition in a cell by an engineered Himar-dCas9 system for the first time.

## DISCUSSION

In this study, we showed the successful engineering of a targeted transposase for *in vitro* and *in vivo* site-specific transposon insertion. The Himar-dCas9 fusion protein, consisting of a hyperactive Himar1C9 transposase and the *S. pyogenes* dCas9, functions to mediate site-specific transposition into a targeted genetic locus. We characterized the activity of Himar-dCas9 *in vitro* across different protein and DNA concentrations, as well as various reaction conditions, demonstrating that the site-specific transposition activity is robust. We further showed that transposition is dependent on the gRNA orientation relative to the target TA insertion site and the surrounding target DNA sequence. In *in vivo* studies of Himar-dCas9 in *E. coli*, we showed that gRNA_1 successfully targeted transposition to a single locus on a medium copy-number target plasmid in more than 80% of detected transposition events.

Comparing the results of the *in vitro* and *in vivo* studies, there were persistent off-target transposition events in both types of studies, while gRNA strand complementarity appears to affect site-specific transposition *in vivo* but not *in vitro*. We attribute the off-target transposition to constitutive activity of Himar1C9 domains within the Himar-dCas9 fusion protein, even in transposase dimers which are not bound to the gRNA-targeted site. The difference in transposition targeting by gRNA_1 and gRNA_4, which both resulted in targeted transposition *in vitro*, may be explained by the presence of other DNA-interacting elements *in vivo*. As discussed above, gRNA_1-guided Himar-dCas9 bound the non-template strand of the GFP gene, sterically hindering RNA polymerase from traversing downstream to its target TA site, while gRNA_4-guided Himar-dCas9 did not hinder RNA polymerase from accessing its upstream TA site, unwinding the target DNA, and blocking transposition. This difference in relative accessibility of the targeted TA site by Himar-dCas9 versus RNA polymerase may explain the strand-dependence of *in vivo* transposition targeting by gRNAs.

Protein engineering of Himar-dCas9 could further improve efficiency of transposition and reduce the frequency of off-target transposition. DNA base editors consisting of a deaminase fused to a dCas9 protein have improved efficiency when two monomers of the deaminase, which operates as a homodimer, are fused to dCas9^35^. Because Himar1 likewise functions as a homodimer, the dimer-dCas9 fusion approach may be useful for targeting active Himar1 dimers to the desired insertion site. Modification of the Himar1 protein by directed evolution or by rational mutagenesis may also improve function. Previous studies have shown that Himar1, like other *mariner* transposases, is rate-limited by the synapsis of transposon ends to the protein dimer^48, 49^. Himar1 mutants with single amino-acid substitutions in the conserved WVPHEL motif have higher rates of transposition than the Himar1C9 mutant, because the allosteric inhibition of transposon synapsis is disrupted^48^. Thus, it is possible that a hyperactive WVPHEL mutation, which removes the natural rate-limiting step of transposon synapsis, combined with a mutation in the dimerization interface to slow down dimerization of Himar1 (normally a fast step), may result in a Himar1 protein that dimerizes more specifically at dCas9-targeted locations and then efficiently catalyzes transposition at those sites. Alternative protein components can also be explored, including other transposases (e.g., Tn5), DNA-binding domains (e.g., Cas homologs), and peptide linkers.

While targeted insertion was demonstrated here for a medium copy-number target plasmid in *E. coli*, additional optimizations may be necessary to improve Himar-dCas9 targeting to a genomic locus and detection of targeted genomic insertion events. Bacterial assays for targeted genomic transposition using counterselectable genomic markers, such as *galK, tolC*, or *sacB*, as targets may be useful for detecting site-specific transposon insertions^50-52^. The use of inducible Himar-dCas9 can improve the differentiation between gene knockout (via targeted transposition) and knockdown (via transcriptional blockade by Himar-dCas9) to improve the sensitivity and selectivity of insertion events. To reduce leakiness of inducible expression systems which may yield unintended off-target transpositions, further improvements could be made to transiently express the Himar-dCas9/gRNA complex through acquisition of a non-replicative vector (e.g. a R6K plasmid) in the target cell by conjugation or transformation. Transient presence of the transposon donor vector would also be a useful feature in genomic targeting studies so that the pool of integrated transposons could be isolated and analyzed by Tn-seq. Optimized Cas-Transposon technologies can serve as a platform to enable a novel modality of site-specific DNA insertion for targeted genome editing.

## ACKNOWLEDGEMENTS

We thank Saeed Tavazoie, Alexandra Ketcham, and Anupama Khare for guidance on transposon sequencing. Eric Greene, Justin Steinfeld, and Chu Jian Ma provided materials and guidance on *in vitro* protein expression and purification. We thank members of the Wang lab for helpful scientific discussions and feedback.

## AUTHOR CONTRIBUTIONS

S.P.C. and H.H.W. designed the study. S.P.C. performed the experiments. S.P.C. and H.H.W. analyzed the data and wrote the manuscript.

## References

1. Esvelt, K.M. & Wang, H.H. Genome-scale engineering for systems and synthetic biology. Molecular systems biology 9, 641 (2013).

2. Andrews, B.J., Proteau, G.A., Beatty, L.G. & Sadowski, P.D. The FLP recombinase of the 2 micron circle DNA of yeast: interaction with its target sequences. Cell 40, 795–803 (1985).

3. Abremski, K. & Hoess, R. Bacteriophage P1 site-specific recombination. Purification and properties of the Cre recombinase protein. The Journal of biological chemistry 259, 1509–1514 (1984).

4. Bolusani, S. et al. Evolution of variants of yeast site-specific recombinase Flp that utilize native genomic sequences as recombination target sites. Nucleic acids research 34, 5259–5269 (2006).

5. Buchholz, F. & Stewart, A.F. Alteration of Cre recombinase site specificity by substrate-linked protein evolution. Nature biotechnology 19, 1047–1052 (2001).

6. Cong, L. et al. Multiplex genome engineering using CRISPR/Cas systems. Science 339, 819–823 (2013).

7. Jinek, M. et al. A programmable dual-RNA-guided DNA endonuclease in adaptive bacterial immunity. Science 337, 816–821 (2012).

8. Urnov, F.D., Rebar, E.J., Holmes, M.C., Zhang, H.S. & Gregory, P.D. Genome editing with engineered zinc finger nucleases. Nature reviews. Genetics 11, 636–646 (2010).

9. Joung, J.K. & Sander, J.D. TALENs: a widely applicable technology for targeted genome editing. Nat Rev Mol Cell Biol 14, 49–55 (2013).

10. Kowalczykowski, S.C. An Overview of the Molecular Mechanisms of Recombinational DNA Repair. Cold Spring Harb Perspect Biol 7 (2015).

11. Munoz-Lopez, M. & Garcia-Perez, J.L. DNA transposons: nature and applications in genomics. Curr Genomics 11, 115–128 (2010).

12. Curcio, M.J. & Derbyshire, K.M. The outs and ins of transposition: from mu to kangaroo. Nat Rev Mol Cell Biol 4, 865–877 (2003).

13. Lampe, D.J., Churchill, M.E. & Robertson, H.M. A purified mariner transposase is sufficient to mediate transposition in vitro. The EMBO journal 15, 5470–5479 (1996).

14. Richardson, J.M. et al. Mechanism of Mos1 transposition: insights from structural analysis. The EMBO journal 25, 1324–1334 (2006).

15. Richardson, J.M., Colloms, S.D., Finnegan, D.J. & Walkinshaw, M.D. Molecular architecture of the Mos1 paired-end complex: the structural basis of DNA transposition in a eukaryote. Cell 138, 1096–1108 (2009).

16. Claeys Bouuaert, C., Lipkow, K., Andrews, S.S., Liu, D. & Chalmers, R. The autoregulation of a eukaryotic DNA transposon. eLife 2, e00668 (2013).

17. van Opijnen, T. & Camilli, A. Transposon insertion sequencing: a new tool for systems-level analysis of microorganisms. Nature reviews. Microbiology 11, 435–442 (2013).

18. Zhang, L., Sankar, U., Lampe, D.J., Robertson, H.M. & Graham, F.L. The Himar1 mariner transposase cloned in a recombinant adenovirus vector is functional in mammalian cells. Nucleic acids research 26, 3687–3693 (1998).

19. Lampe, D.J., Grant, T.E. & Robertson, H.M. Factors affecting transposition of the Himar1 mariner transposon in vitro. Genetics 149, 179–187 (1998).

20. Lampe, D.J., Akerley, B.J., Rubin, E.J., Mekalanos, J.J. & Robertson, H.M. Hyperactive transposase mutants of the Himar1 mariner transposon. Proceedings of the National Academy of Sciences of the United States of America 96, 11428–11433 (1999).

21. Goodman, A.L. et al. Identifying genetic determinants needed to establish a human gut symbiont in its habitat. Cell Host Microbe 6, 279–289 (2009).

22. van Opijnen, T., Bodi, K.L. & Camilli, A. Tn-seq: high-throughput parallel sequencing for fitness and genetic interaction studies in microorganisms. Nature methods 6, 767–772 (2009).

23. Zhang, J.K., Pritchett, M.A., Lampe, D.J., Robertson, H.M. & Metcalf, W.W. In vivo transposon mutagenesis of the methanogenic archaeon Methanosarcina acetivorans C2A using a modified version of the insect mariner-family transposable element Himar1. Proceedings of the National Academy of Sciences of the United States of America 97, 9665–9670 (2000).

24. Maragathavally, K.J., Kaminski, J.M. & Coates, C.J. Chimeric Mos1 and piggyBac transposases result in site-directed integration. FASEB J 20, 1880–1882 (2006).

25. Owens, J.B. et al. Chimeric piggyBac transposases for genomic targeting in human cells. Nucleic acids research 40, 6978–6991 (2012).

26. Owens, J.B. et al. Transcription activator like effector (TALE)-directed piggyBac transposition in human cells. Nucleic acids research 41, 9197–9207 (2013).

27. Luo, W. et al. Comparative analysis of chimeric ZFP-, TALE- and Cas9-piggyBac transposases for integration into a single locus in human cells. Nucleic acids research 45, 8411–8422 (2017).

28. Feng, X., Bednarz, A.L. & Colloms, S.D. Precise targeted integration by a chimaeric transposase zinc-finger fusion protein. Nucleic acids research 38, 1204–1216 (2010).

29. Morero, N.R. et al. Targeting IS608 transposon integration to highly specific sequences by structure-based transposon engineering. Nucleic acids research 46, 4152–4163 (2018).

30. Adli, M. The CRISPR tool kit for genome editing and beyond. Nature communications 9, 1911 (2018).

31. Qi, L.S. et al. Repurposing CRISPR as an RNA-guided platform for sequence-specific control of gene expression. Cell 152, 1173–1183 (2013).

32. Gilbert, L.A. et al. CRISPR-mediated modular RNA-guided regulation of transcription in eukaryotes. Cell 154, 442–451 (2013).

33. Guilinger, J.P., Thompson, D.B. & Liu, D.R. Fusion of catalytically inactive Cas9 to FokI nuclease improves the specificity of genome modification. Nature biotechnology 32, 577–582 (2014).

34. Tsai, S.Q. et al. Dimeric CRISPR RNA-guided FokI nucleases for highly specific genome editing. Nature biotechnology 32, 569–576 (2014).

35. Gaudelli, N.M. et al. Programmable base editing of A*T to G*C in genomic DNA without DNA cleavage. Nature 551, 464–471 (2017).

36. Komor, A.C., Kim, Y.B., Packer, M.S., Zuris, J.A. & Liu, D.R. Programmable editing of a target base in genomic DNA without double-stranded DNA cleavage. Nature 533, 420–424 (2016).

37. Chaikind, B., Bessen, J.L., Thompson, D.B., Hu, J.H. & Liu, D.R. A programmable Cas9-serine recombinase fusion protein that operates on DNA sequences in mammalian cells. Nucleic acids research 44, 9758–9770 (2016).

38. Pickens, L.B., Tang, Y. & Chooi, Y.H. Metabolic engineering for the production of natural products. Annu Rev Chem Biomol Eng 2, 211–236 (2011).

39. Esvelt, K.M., Smidler, A.L., Catteruccia, F. & Church, G.M. Concerning RNA-guided gene drives for the alteration of wild populations. eLife 3 (2014).

40. Goryshin, I.Y., Miller, J.A., Kil, Y.V., Lanzov, V.A. & Reznikoff, W.S. Tn5/IS50 target recognition. Proceedings of the National Academy of Sciences of the United States of America 95, 10716–10721 (1998).

41. Ronda, C., Chen, S.P., Cabral, V., Yaung, S.J. & Wang, H.H. Metagenomic engineering of the mammalian gut microbiome in situ. Nature methods 16, 167–170 (2019).

42. Rohland, N. & Reich, D. Cost-effective, high-throughput DNA sequencing libraries for multiplexed target capture. Genome research 22, 939–946 (2012).

43. Langmead, B. & Salzberg, S.L. Fast gapped-read alignment with Bowtie 2. Nature methods 9, 357–359 (2012).

44. Sundaresan, R., Parameshwaran, H.P., Yogesha, S.D., Keilbarth, M.W. & Rajan, R. RNA-Independent DNA Cleavage Activities of Cas9 and Cas12a. Cell reports 21, 3728–3739 (2017).

45. Vigdal, T.J., Kaufman, C.D., Izsvak, Z., Voytas, D.F. & Ivics, Z. Common physical properties of DNA affecting target site selection of sleeping beauty and other Tc1/mariner transposable elements. Journal of molecular biology 323, 441–452 (2002).

46. Trubitsyna, M., Morris, E.R., Finnegan, D.J. & Richardson, J.M. Biochemical characterization and comparison of two closely related active mariner transposases. Biochemistry 53, 682–689 (2014).

47. Milo, R., Jorgensen, P., Moran, U., Weber, G. & Springer, M. BioNumbers--the database of key numbers in molecular and cell biology. Nucleic acids research 38, D750–753 (2010).

48. Lampe, D.J. Bacterial genetic methods to explore the biology of mariner transposons. Genetica 138, 499–508 (2010).

49. Liu, D. & Chalmers, R. Hyperactive mariner transposons are created by mutations that disrupt allosterism and increase the rate of transposon end synapsis. Nucleic acids research 42, 2637–2645 (2014).

50. Warming, S., Costantino, N., Court, D.L., Jenkins, N.A. & Copeland, N.G. Simple and highly efficient BAC recombineering using galK selection. Nucleic acids research 33, e36 (2005).

51. Li, X.T., Thomason, L.C., Sawitzke, J.A., Costantino, N. & Court, D.L. Positive and negative selection using the tetA-sacB cassette: recombineering and P1 transduction in Escherichia coli. Nucleic acids research 41, e204 (2013).

52. DeVito, J.A. Recombineering with tolC as a selectable/counter-selectable marker: remodeling the rRNA operons of Escherichia coli. Nucleic acids research 36, e4 (2008).

