## Supplementary Materials for "An Engineered Cas-Transposon System for Programmable and Precise DNA Transpositions"

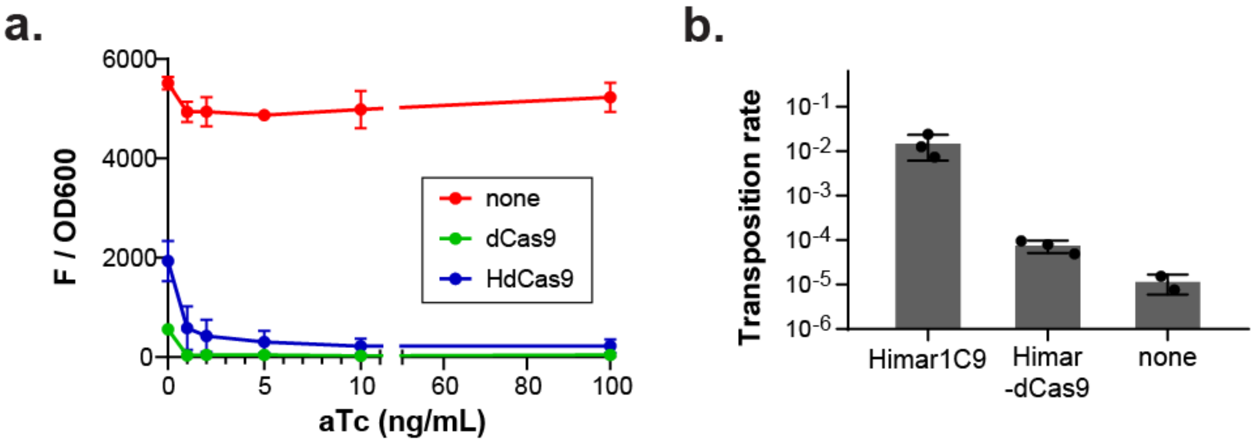


**Supplementary Figure S1. Himar1C9-dCas9 (Himar-dCas9) fusion protein retains DNA binding and transposition functionalities. (a)** dCas9 and Himar-dCas9 were expressed in MG1655 *galK::mCherry-specR E. coli* with gRNAs 5 and 16. Protein expression was induced with aTc (0-100 ng/mL); n=3 for each condition. Both proteins decreased mCherry expression compared with the parent strain, indicating that the Himar-dCas9 fusion protein bound to the mCherry gene specified by the gRNAs and blocked transcription. **(b)** The transposition rates of Himar1C9 and Himar-dCas9 (without gRNA) were measured in an *E. coli* conjugation assay (n=3 for transposases, n=2 for control). Both Himar1C9 and Himar-dCas9 mediated transposition at higher rates than the no-transposase control.

Error bars indicate standard deviation.

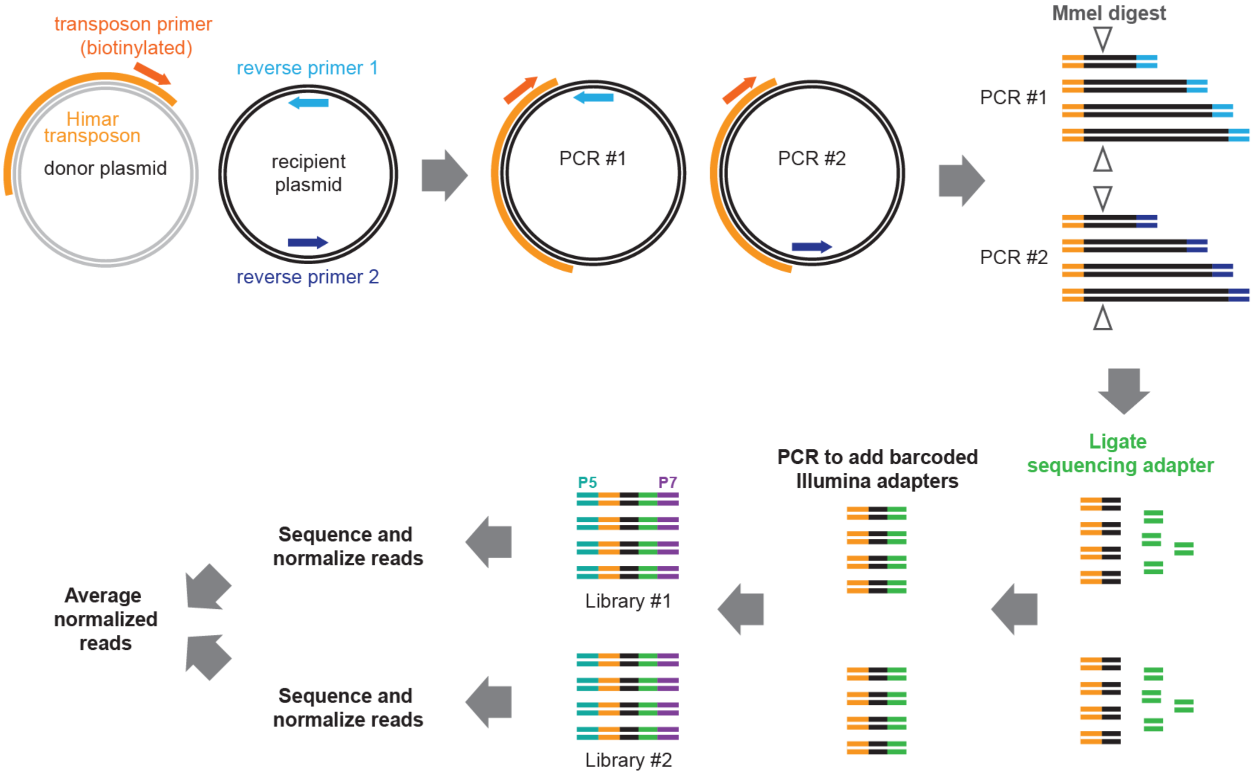


**Supplementary Figure S2. Workflow for transposon sequencing library preparation from *in vitro* transposition reactions.** To selectively isolate transposons that had become integrated into recipient plasmid for sequencing, we performed PCRs using a biotinylated primer complementing the transposon end and reverse primers complementing the recipient plasmid. Two PCRs using reverse primers on opposite sides of the recipient plasmid were performed to account for PCR size bias during amplification of transposon junction products. PCR products were isolated using streptavidin beads and digested with MmeI to isolate transposon ends with a ~17bp overhang. A sequencing adapter was ligated, and the DNA was PCR-amplified to add barcoded Illumina adapters. The resulting libraries from each PCR were sequenced independently and normalized for total reads, and the normalized libraries were averaged to obtain transposon insertion frequencies into each locus on the plasmid.

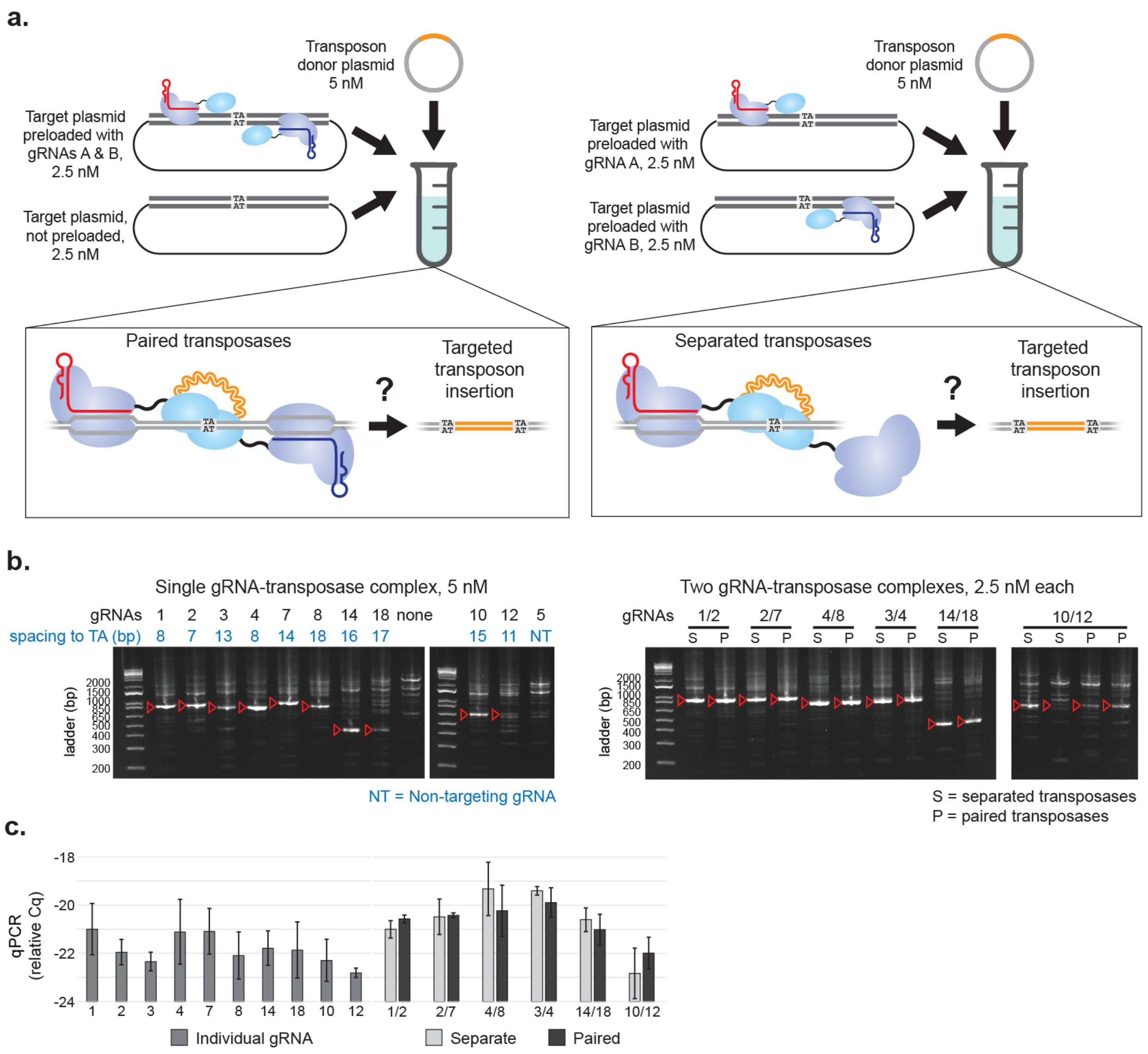


**Supplementary Figure S3. *In vitro* assay to analyze transposition by Himar-dCas9 with 2 gRNAs. (a)** *In vitro* reactions containing 2 gRNAs were set up in 2 configurations to determine whether paired Himar-dCas9 proteins bound at the same TA site would improve transposase dimerization and activity, compared with Himar-dCas9 proteins all bound individually to target plasmids. Himar-dCas9 was first incubated with either gRNA A (red) or gRNA B (blue), and then the Himar-dCas9/gRNA complexes were preloaded onto target plasmids as pairs (left) or as single complexes (right). Preloaded target plasmid-Himar-dCas9/gRNA complexes were then mixed with transposon donor plasmids. The total final concentration of each protein-gRNA complex was 2.5 nM, and final concentrations of donor and target DNAs were 5 nM. **(b)** PCR analysis of transposition by Himar-dCas9 with a single gRNA (left) or Himar-dCas9 with 2 gRNAs (right), preloaded in separated (S) or paired configurations (P). Red arrows indicate expected PCR products for each reaction. **(c)** qPCR analysis of transposition by Himar-dCas9 with a single gRNA, Himar-dCas9 with 2 gRNAs (in a separated configuration), and Himar-dCas9 with 2 gRNAs (in a paired configuration). n = 2-6 reactions per condition; error bars indicate standard deviation.

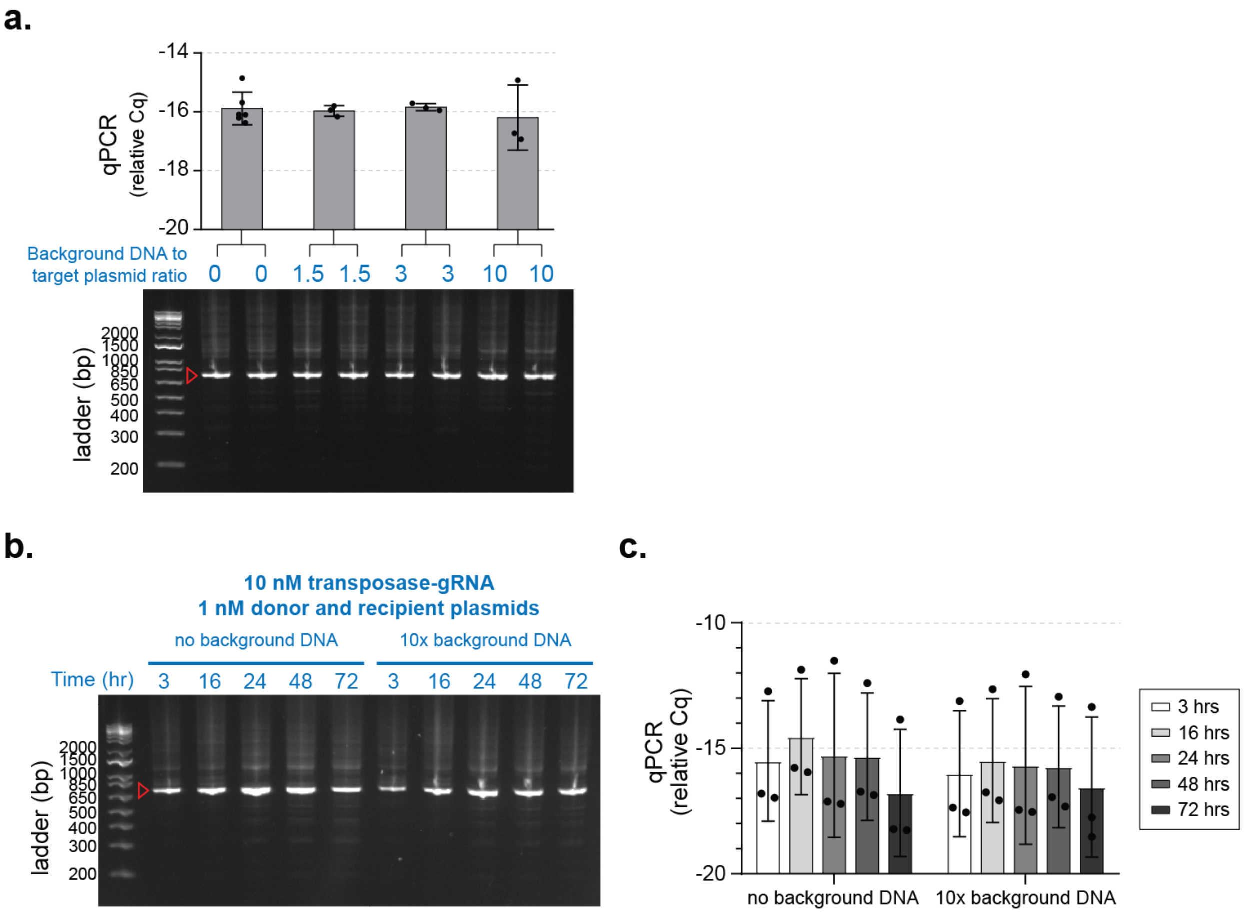


**Supplementary Figure S4. Himar-dCas9 performs *in vitro* site-specific transposition in the presence of background DNA. (a)** PCR analysis of transposition reactions (n=3-6) with varying levels of background *E. coli* genomic DNA. Reactions were performed for 3 hours at 30 °C with 1 nM of target plasmid DNA, 1 nM of donor plasmid DNA, and 10 nM HdCas-gRNA_4 complex. Ratios of background to target plasmid DNA are by mass. **(b)** PCR analysis of transposition reactions (n=3) performed for different lengths of time, in the presence or absence of background non-specific DNA. Reactions were performed at 37 °C with 1 nM of recipient plasmid DNA, 1 nM of donor plasmid DNA, and 10 nM HdCas-gRNA_4 complex. Background *E. coli* genomic DNA was present at 10x the mass of recipient plasmid DNA. **(c)** qPCR measurement of transposition efficiency in reactions shown in panel (b). n=3 for each reaction condition.

Error bars indicate standard deviation.

**Table 1. Plasmids used in this study.**

| Plasmid | Origin of replication | Size (bp) | Selection | Features | Purpose |
|---|---|---|---|---|---|
| pET-Himar-dCas9 | ROP | 10864 | carb | 6xHis tag, T7 promoter | HdCas9 protein purification |
| pGT-B1 | pBBR1 | 6235 | carb | constitutive sfGFP gene | target plasmid for in vitro assays |
| pHimar6 | R6K | 3394 | kan | Himar transposon with chlor resistance cassette, RP4 oriT | Himar transposon donor plasmid for in vitro and E. coli in vivo assays |
| pTarget | ColE1 | 3237 | spec | constitutive sfGFP gene | target plasmid for E. coli in vivo assays |
| pHimar1C9 | p15A | 3846 | carb | Himar1C9 on tet-inducible promoter | bacterial expression vector for Himar1C9 |
| pHdCas9-gRNA1 | p15A | 8200 | carb | Himar-dCas9 on tet-inducible promoter, constitutively expressed gRNA_1 | bacterial expression vector for Himar-dCas9 and gRNA_1 |
| pHdCas9-gRNA4 | p15A | 8200 | carb | Himar-dCas9 on tet-inducible promoter, constitutively expressed gRNA_4 | bacterial expression vector for Himar-dCas9 and gRNA_4 |
| pHdCas9-gRNA5 | p15A | 8200 | carb | Himar-dCas9 on tet-inducible promoter, constitutively expressed gRNA_5 | bacterial expression vector for Himar-dCas9 and gRNA_5 |
| pHdCas9 | p15A | 7738 | carb | Himar-dCas9 on tet-inducible promoter | bacterial expression vector for Himar-dCas9 |
| pdCas9-carb | p15A | 6847 | carb | dCas9 on tet-inducible promoter | bacterial expression vector for Himar-dCas9 |
| pHdCas9-gRNA5-gRNA16 | p15A | 8191 | chlor | Himar-dCas9 on tet-inducible promoter, constitutively expressed gRNA_5 and gRNA_16 | bacterial expression vector for Himar-dCas9, gRNA_5, gRNA_16 |
| pdCas9-gRNA5-gRNA16 | p15A | 7099 | chlor | dCas9 on tet-inducible promoter, constitutively expressed gRNA_5 and gRNA_16 | bacterial expression vector for dCas9, gRNA_5, gRNA_16 |

**Table 2. gRNA sequences used in this study.**

| gRNA name | Sequence | Target gene | Target strand (T/N) | Spacing to TA site (bp) |
|---|---|---|---|---|
| gRNA_1 | GTCGTTACCAGAGTCGGCCA | sfGFP | N | 8 |
| gRNA_2 | TCAGTGCTTTGCTCGTTATC | sfGFP | T | 7 |
| gRNA_3 | CGTTCCTGCACATAGCCTTC | sfGFP | N | 13 |
| gRNA_4 | CGGCACGTACAAAACGCGTG | sfGFP | T | 8 |
| gRNA_5 | GTCGGCGGGGTGCTTCACGT | mCherry | N | 10 |
| gRNA_7 | ACCAGAGTCGGCCAAGGTAC | sfGFP | N | 14 |
| gRNA_8 | CTGCACATAGCCTTCCGGCA | sfGFP | N | 18 |
| gRNA_9 | CAATGCCTTTCAGCTCAATG | sfGFP | N | 5 |
| gRNA_10 | CAGCTCAATGCGGTTTACCA | sfGFP | N | 15 |
| gRNA_11 | GTAAACCGCATTGAGCTGAA | sfGFP | T | 6 |
| gRNA_12 | CAATATCCTGGGCCATAAGC | sfGFP | T | 11 |
| gRNA_13 | AGAACAGGACCATCACCGAT | sfGFP | N | 17 |
| gRNA_14 | GTGCTCAGATAGTGATTGTC | sfGFP | N | 16 |
| gRNA_15 | GAACTGGATGGTGATGTCAA | sfGFP | T | 9 |
| gRNA_16 | CCTTCCCCGAGGGCTTCAAG | mCherry | T | 12 |
| gRNA_18 | ACGCGATCACATGGTTCTGC | sfGFP | T | 17 |

T indicates that the gRNA is complementary to the Template strand of the gene, while N indicates that the gRNA complements the Non-template strand. gRNAs that target the same TA insertion site are labeled with the same color. gRNAs 11, 13, and 15 all target different sites uniquely.

**Table 3. Oligonucleotides used in this study.**

| Name | Sequence (5' -> 3') | Target | Tm (°C) | Function |
|---|---|---|---|---|
| p433 | CGCTTACAATTTCCATTC GCCATTC | pGT-B1 | 67 | qPCR for Himar transposon-pGT-B1 junction |
| p415 | CCCTGCAAAGCCCCTCTT TACG | pHimar6 transposon | 71 | qPCR for Himar transposon-pGT-B1 junction |
| p828 | CTGCGCAACCCAAGTGCT AC | pGT-B1 | 70 | Control qPCR for pGT-B1 |
| p829 | CAGTCCAGAGAAATCGGC ATTCA | pGT-B1 | 67 | Control qPCR for pGT-B1 |
| p923 | Biotin/GCCATAAACTG CCAGGCATCAA | pHimar6 transposon | 68 | In vitro transposon sequencing library preparation |
| p922 | CCTTCTTGCGCATCTCAC G | pGT-B1 | 67 | In vitro transposon sequencing library preparation |
| Adapter_T | Phosphate/AGATCGGA AGAGCACACGTCTG AACTCCAGTCAC | | | Anneal to make Y-shapted adapter for Tn-seq library prep |
| Adapter_B | GTCTCGTGGGCTCGGGCT CTTCCGATCT*N*N | | | Anneal to make Y-shapted adapter for Tn-seq library prep |
| p790 | AATGATACGGCGACCACC GAGATCTacacTAGATCG CCGCCagaccggggactt atcatccaacctgt | Himar transposon IR | 73 | Add barcode & P5 sequence to Himar transposon ends for Illumina sequencing |
| p791 | AATGATACGGCGACCACC GAGATCTacacCTCTCTA TCGCCagaccggggactt atcatccaacctgt | Himar transposon IR | 73 | Add barcode & P5 sequence to Himar transposon ends for Illumina sequencing |
| p792 | AATGATACGGCGACCACC GAGATCTacacTATCCTC TCGCCagaccggggactt atcatccaacctgt | Himar transposon IR | 73 | Add barcode & P5 sequence to Himar transposon ends for Illumina sequencing |
| p793 | AATGATACGGCGACCACC GAGATCTacacAGAGTAG ACGCCagaccggggactt atcatccaacctgt | Himar transposon IR | 73 | Add barcode & P5 sequence to Himar transposon ends for Illumina sequencing |
| p794 | AATGATACGGCGACCACC GAGATCTacacGTAAGGA GCGCCagaccggggactt atcatccaacctgt | Himar transposon IR | 73 | Add barcode & P5 sequence to Himar transposon ends for Illumina sequencing |
| p795 | AATGATACGGCGACCACC GAGATCTacacACTGCAT ACGCCagaccggggactt atcatccaacctgt | Himar transposon IR | 73 | Add barcode & P5 sequence to Himar transposon ends for Illumina sequencing |
| p712 | CGCCagaccggggactta tcatccaacctgt | Himar transposon IR | 67 | Read 1 primer for Illumina sequencing |
| p713 | CGGAAGAGCCCGAGCCCA CGAGAC | Himar sequencing library | 67 | Index 1 primer for Illumina sequencing |
| p898 | TTTGAGTGAGCTGATACC GCTC | ColE1 oriR | 67 | qPCR for Himar transposon-plasmid junctions in pTarget plasmid |
| p899 | GAGCGGTATCAGCTCACT CAAA | ColE1 oriR | 67 | Control qPCR for pTarget |
| p900 | TCCCTTAACGTGAGTTTT CGTTCC | ColE1 oriR | 67 | Control qPCR for pTarget |
